# 3D Deconvolution Processing for STEM Cryo-Tomography

**DOI:** 10.1101/2020.08.26.267732

**Authors:** Barnali Waugh, Sharon G. Wolf, Deborah Fass, Eric Branlund, Zvi Kam, John Sedat, Michael Elbaum

**Affiliations:** Dept of Chemical and Biological Physics, Weizmann Institute of Science, Rehovot 7610001, Israel; Chemical Research Support Dept, Weizmann Institute of Science, Rehovot 7610001, Israel; Dept of Structural Biology, Weizmann Institute of Science, Rehovot 7610001, Israel; Dept of Molecular Cell Biology, Weizmann Institute of Science, Rehovot 7610001, Israel; Department of Biochemistry and Biophysics, University of California, San Francisco, CA 94158 USA

## Abstract

The complex environment of biological cells and tissues has motivated development of three dimensional imaging in both light and electron microscopies. To this end, one of the primary tools in fluorescence microscopy is that of computational deconvolution. Wide-field fluorescence images are often corrupted by haze due to out-of-focus light, i.e., to cross-talk between different object planes as represented in the 3D image. Using prior understanding of the image formation mechanism, it is possible to suppress the cross-talk and reassign the unfocused light to its proper source *post facto*. Electron tomography based on tilted projections also exhibits a cross-talk between distant planes due to the discrete angular sampling and limited tilt range. By use of a suitably synthesized 3D point spread function, we show here that deconvolution leads to similar improvements in volume data reconstructed from cryo-scanning transmission electron tomography (CSTET), namely a dramatic in-plane noise reduction and improved representation of features in the axial dimension. Contrast enhancement is demonstrated first with colloidal gold particles, and then in representative cryo-tomograms of intact cells. Deconvolution of CSTET data collected from the periphery of an intact nucleus revealed partially condensed, extended structures in interphase chromatin.

**Significance statement:** Electron tomography is used to reveal the structure of cells in three dimensions. The combination with cryogenic fixation provides a snapshot in time of the living state. However, cryo-tomography normally requires very thin specimens due to image formation by conventional phase contrast transmission electron microscopy (TEM). The thickness constraint can be relaxed considerably by scanning TEM (STEM), yet three-dimensional (3D) reconstruction is still subject to artifacts inherent in the collection of data by tilted projections. We show here that deconvolution algorithms developed for fluorescence microscopy can suppress these artifacts, resulting in significant contrast enhancement. The method is demonstrated by cellular tomography of complex membrane structures, and by segmentation of chromatin into distinct, contiguous domains of heterochromatin and euchromatin at high and low density, respectively.

## Introduction

Electron tomography (ET) is the premier technique for visualization of cellular ultrastructure in three dimensions. ET actually encompasses a number of different experimental approaches (1–3). These include serial sections imaged by transmission electron microscopy (TEM) and “serial surface” views produced by scanning electron microscopy (SEM) combined with serial or iterative microtomy. A 3D image is produced by aligning and combining the sections. Most commonly, ET is performed by rotating a specimen around an axis while a series of projection images is recorded by wide-field TEM. A 3D image can then be reconstructed from the projections by standard algorithms (4, 5). ET is routinely performed on plastic embedded cell or tissue specimens that have been stained with heavy metals for contrast enhancement. ET on unstained hydrated specimens at cryogenic temperatures can provide even molecular detail (6, 7).

Cryogenic vitrification offers potentially the most faithful preservation of biological cells and other aqueous-based materials (8). Imaging without the benefit of contrast-enhancing stains is highly challenging, however. The combination of weak electron scattering and strict exposure limits to avoid specimen damage results in noisy images. In addition, certain systematic artifacts are inherent to reconstructions from projection tilt series. While in specific cases a cylindrical geometry provides for full rotation around the tilt axis (9, 10), the more typical flat slab-geometry specimens can be tilted only within a certain restricted range. This artifact is known as the “missing wedge”; the missing projections impair the reconstruction. Acquisition of projection images at a limited number of discrete angles also results in missing information for reconstruction (11, 12). Both limitations lead to “ghost” contrast that emanates from sharp features and projects into neighboring planes.

Ghost artifacts and the smearing of axial contrast in electron tomography recall out-of-focus blur in fluorescent light microscopy. Fluorescence deconvolution algorithms have reached a degree of maturity, yet advances are also underway (13). In particular, entropy regularization makes a significant improvement in noise suppression (14). Assuming that the 3D image of a point source, i.e., the point spread function (PSF), is known, one might similarly reassign the reconstruction artifacts of electron tomography back to the plane of origin by 3D image deconvolution. In this work we ask whether algorithms intended for optical deconvolution may serve to enhance 3D reconstruction by electron cryo-tomography.

At first sight, the very different configurations of fluorescence and electron microscopies make such an application unlikely. Fluorescence emission behaves as a source of light; its intensity is (within limits) quantitatively proportional to the local fluorophore density in the specimen. Transmission electron microscopy detects electron scattering by the projected electrostatic potential, which is (very) approximately proportional to electron density; the scattering contrast between water and organic materials is inherently low. Fluorescence imaging is typically performed with objective lenses of high numerical aperture with a cylindrically symmetric PSF covering as much as 70 degrees in collection semi-angle. For cryo-electron tomography, projection images are normally recorded one by one using parallel illumination and phase contrast in wide field TEM. Fluorescence contrast is unipolar – bright on a dark background. Phase images are interferometric by nature; they contain areas both brighter and darker than background due to constructive and destructive interference. Reliance on phase contrast also introduces a number of complications specifically for thick specimens. These originate from inelastic scattering and chromatic aberrations, on one hand, and from violation of the weak phase object approximation on the other (15).

Tomography by scanning TEM (STEM) was introduced for plastic embedded section (16) and offers an alternative to wide field phase contrast for cryo-tomography (17–20). In STEM, a focused probe is scanned across the specimen while independent detectors record the scattered (dark-field) and unscattered (bright-field) signals. The convergence angle of the illumination cone, as well as the angular ranges collected by the detectors, are under user control (20–22). Under incoherent imaging conditions, contrast depends on differential scattering cross-sections weighted by the densities of constituent atoms and integrated over the detector apertures. For cryo-scanning transmission electron tomography (CSTET), a small convergence angle is often used in order to extend the depth of field (18, 23) while dynamic focusing adjusts the focus as a function of specimen tilt and distance of the scanned probe from the tilt axis. Scanned images recorded over a range of specimen tilts can be reconstructed conventionally by back-projection. Linearity of the image contrast does not depend on the weak phase limit so the projection assumption for tomography is better satisfied by STEM than by TEM for thick specimens (20). In practice, the limiting thickness may exceed one micron.

Deconvolution is based on the image of an ideal point source, the PSF. In STEM, incoherent electron scattering in the specimen acts as a local source, similarly to fluorescence emission in light microscopy. Per image, the PSF is approximated simply by the illumination profile, i.e., a diffraction-limited beam focused on the specimen. Since the back-projection operation is essentially additive, an effective three-dimensional PSF can be synthesized as the sum of individual PSFs tilted to the appropriate angles. Rotation around a single axis produces a fan-like pattern in the plane perpendicular. This pattern becomes the 3D PSF for deconvolution of the reconstructed volume (Fig 1). 3D deconvolution suppresses artifactual projections that produce a speckling in distant reconstructed planes and reduce contrast in depth. The effect was seen most clearly with colloidal gold particles. Applied to cellular data, contrast enhancement by deconvolution enhances the visibility of low-contrast features as well. Most strikingly it allowed for sharp distinctions between high- and low-density chromatin regions inside the cell nucleus.

**Figure 1.**
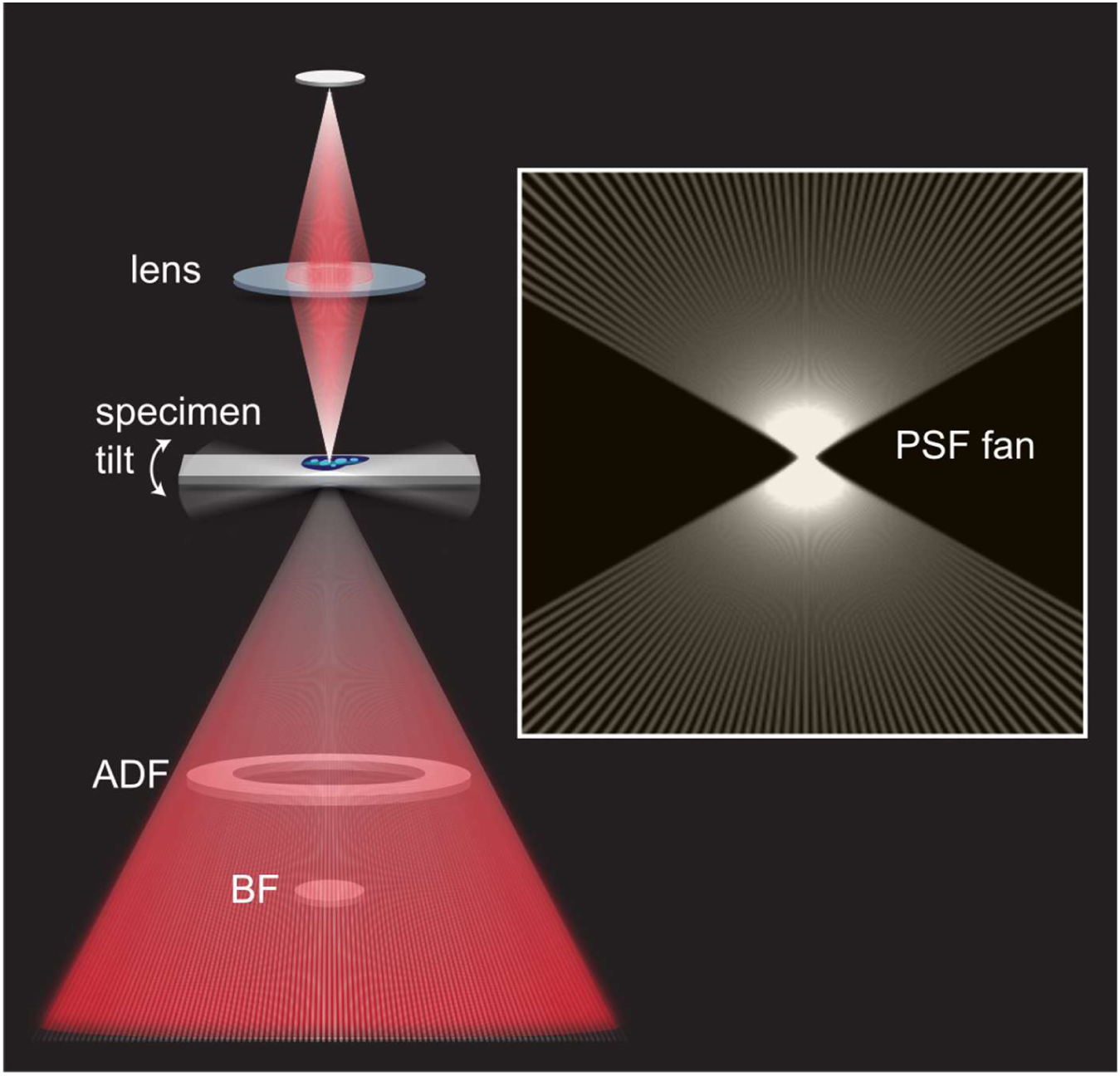
STEM configuration and synthesis of the 3D PSF. A STEM image is formed by scanning a focused electron beam (angles not to scale) across the specimen and collecting the scattered electrons on area detectors. These may include an on-axis bright field (BF) disk and/or a dark field annulus (ADF), each of which integrates the scattered flux over a certain angular range. For tomography, a series of projection images is recorded as the specimen is tilted. The boxed inset shows the construction of the 3D point spread function from a sum of rays representing the illumination profile, tilted to the relevant acquisition angles.

## Methods

### 1. Specimens and microscopy

All data were collected using a Tecnai T-20F S/TEM (FEI, Inc) operating at 200 kV. A condenser C2 aperture of 20 μm was used, resulting in an illumination semi-angle of 2.5 mrad. Tilt series were recorded simultaneously using bright field (BF) and high angle annular dark field (HAADF) detectors (Gatan, Inc. model 807 and Fischione, Inc. model 3000, respectively) from −60° to 60° with an increment of 2°. Image series were aligned in IMOD (24) and reconstructed by weighted back-projection in tomo3d (25).

Gold colloid: A suspension of homemade 6 nm diameter gold colloid was applied to Quantifoil grids (0.6/1R), which were then blotted and plunged to liquid ethane in a Leica EM-GP plunger. Data were acquired at 1.43 nm/pixel. A local region of interest was selected and aligned in IMOD using the gold itself as fiducial markers (24). 3D reconstruction of the HAADF data was generated by weighted back projection (WBP).

Cells: WI-38 human lung fibroblast cells were cultured in full MEM media with 15% fetal calf serum at 37 °C in 5% CO2. Cells were plated on gold Quantifoil R3.5/1 grids and grown for three days. For the purposes of a separate study (26), the sample used for Figs 3-5 was treated with 1 μM oligomycin to suppress mitochondrial respiration. The sample used in Fig 6 was untreated. Grids were blotted and plunged as above with addition of 15 nm diameter gold beads as fiducial markers (27). Data were acquired at 1.68 nm/pixel. BF data were analyzed due to superior resolution for thick specimens (20), with contrast inverted to match the PSF.

### 2. Simulation of single- and multi-probe (fan) PSF

A single probe profile was simulated at the data acquisition condition (refractive index 1.0, numerical aperture 0.0025, wavelength 0.025 Å) by the Diffraction PSF 3D plugin implemented in ImageJ (28). Pixel sampling was chosen to match the original acquisition, or a 2x binned version in Figure 6. A macro in ImageJ was developed to rotate the PSF to the refined projection tilts (.tlt file) generated after final alignment of projections in IMOD, and to sum the corresponding probes to generate the 3D PSF fan. Dimensions were chosen such that the center of the PSF fan coincided with the center of the volume to be deconvolved. All further processing was performed using PRIISM software (29).

### 3. Deconvolution of selected regions of the 3D reconstructions

A specific region of the tomogram was first selected, and the reconstructed volume was deconvolved using the simulated PSF by one of four techniques:

(i)Wiener deconvolution using Iterative Deconvolve 3D (30) in ImageJ.

Wiener deconvolution attempts to recover an approximation of the original specimen distribution from its image as degraded by the instrument, i.e., the point spread function. The parameter gamma was varied from 0 to 0.1, without iteration in order to apply filters only.

(ii)Iterative Deconvolve 3D in ImageJ (30)

The algorithm is an iterative least square solver with positivity constraint. For our calculations parameter gamma was varied from 0.0001 to 0.1; maximum number of iterations was varied from 1 to 50. Calculation was terminated if mean δ < 0.01.

(iii)Deconvolution Lab 2 (31)

Deconvolution Lab 2 provides a java environment (or ImageJ plugin) for deconvolution by a variety of algorithms. Landweber and Richardson-Lucy methods were employed.

(iv)ER-Decon in PRIISM (32)

The regularization implemented in ER-Decon II, a program operating in the PRIISM environment (29), is an entropy based algorithm with positivity constraint (14). First, an optical transfer function (OTF) was generated by 3D Fourier transform of the PSF fan, and then the core2decon algorithm was run to obtain the deconvolution. The standard regularization algorithm with positivity constraint was used throughout. Missing cone weight was disabled. Smoothing and non-linearity parameters were varied as described. Maximum number of iteration was typically set to 50; the algorithm is stable so that more iterations produce only better results (with diminishing returns) at the expense of computation time. Other parameters were left at default values. Full details of the protocol are provided in Supporting Information. Figures were prepared using UCSF Chimera (33).

## Results

Colloidal gold particles provide a convenient first sample to test and calibrate the deconvolution methods. They scatter strongly but lack internal features. Fig 2A shows a reconstruction of 6 nm diameter gold beads embedded in vitreous ice. Starting from the acquired tilt series, images were aligned in IMOD, and the reconstruction was computed using weighted back projection. Orthogonal sections through the reconstructed volume are displayed, with the Z axis representing the beam direction. Fans of rays projecting from each bead are prominently seen in the XZ view. The representation of a featureless bead in the reconstruction is essentially the measured 3D PSF, to be compared with the idealized version seen in Fig 1. Note that IMOD refines numerous parameters as part of the alignment procedure, including the tilt angles. The ray directions in the reconstruction correspond visibly to the refined tilt angles used to generate the back projection, rather than the nominal angles set at the microscope control. The bright rays appear as spurious streaks in the reconstruction seen most clearly in the XZ cuts; their crossing produces a “salt and pepper” noise in the flanking XY planes. In 2D, this structural noise is difficult to distinguish from statistical noise on top of genuine reconstructed density, but it is essentially a ghost of real structure that lies elsewhere in Z. Since the streaks originate with the PSF, it should be possible to suppress them effectively by deconvolution.

**Figure 2.**
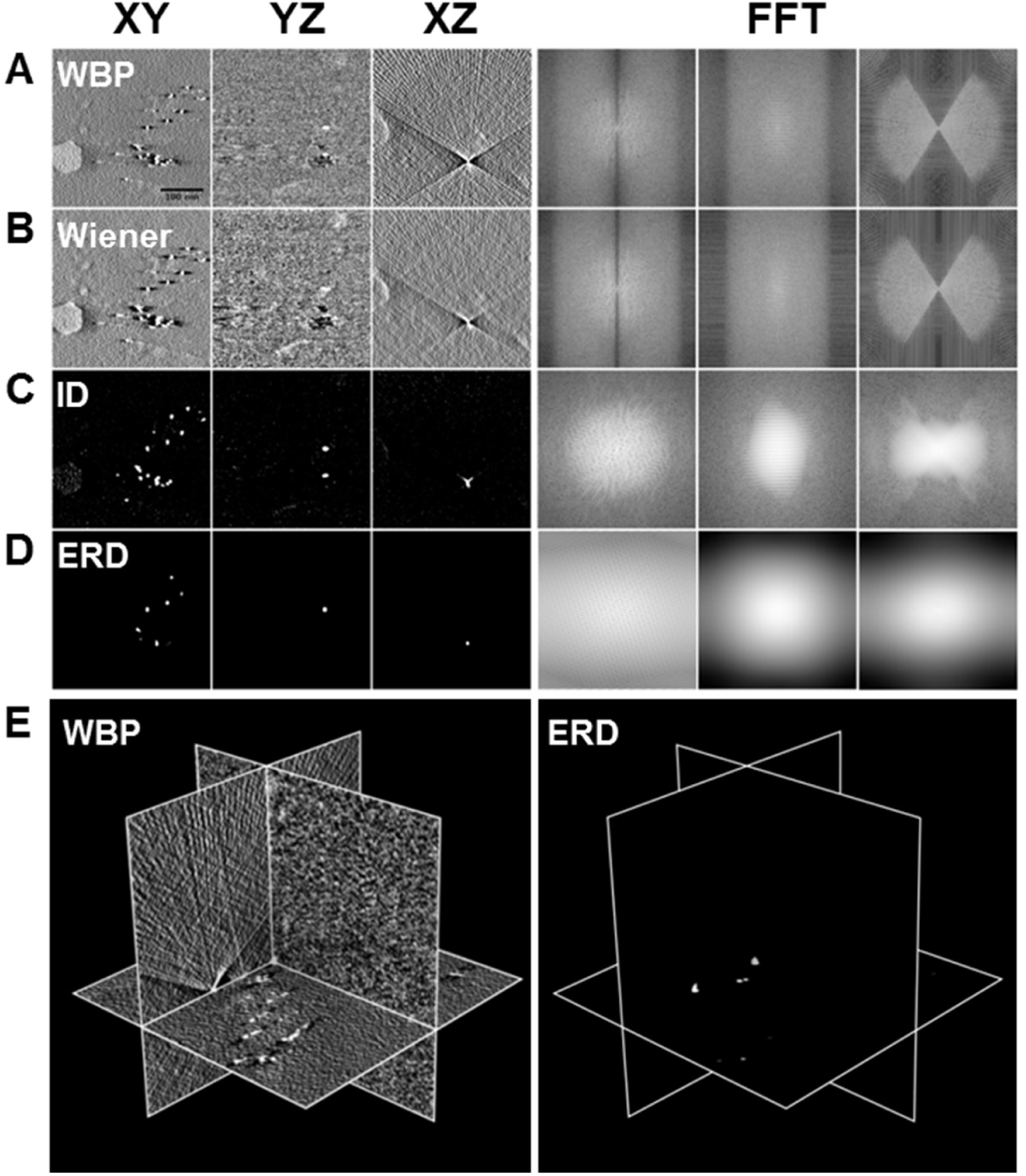
Gold bead data – reconstruction and deconvolution. A) weighted back projection (WBP). Three orthogonal sections (XY, YZ, XZ) passing through the volume of interest. Images to the right (FFT) are Fourier transforms (power spectra, shown in log scale to compress intensities) of the real space planes shown to the left. B) as A for Wiener filtered dataset. C) as A for iterative deconvolution (ID) using ImageJ. D) as A for iterative deconvolution using the ER Decon II algorithm. Parameters for deconvolution were selected visually from a grid shown in Figure S1 of the Supporting Information.

A number of different deconvolution algorithms developed for optical imaging were tested for their effect on the ghost contrast coming from distant planes in the CSTET reconstruction. A 3D PSF was first generated according to the list of refined tilt angles generated in IMOD. Application of a simple Wiener filter had little effect (Fig 2B). After iterative deconvolution (as implemented in the ImageJ plugin Iterative Deconvolve 3D (30)), however, the intensity of the projecting ghosts was dramatically reduced, and only a small hourglass shape remained in the XZ section (Fig 2C). A more recent entropy-regularized algorithm developed for fluorescence imaging (ER-Decon II, (32)) performed still better in restoring the spherical shape of the scattering particle (Fig 2D). The range of operational parameters used in the deconvolution was explored visually in a test grid, as shown in Figure S1 of the Supporting Information. The Landweber and Richardson-Lucy algorithms (implemented in Deconvolution Lab 2 (31)) also suppressed the projection of ghost contrast into neighboring planes (Figure S2 of the SI).

It is instructive to examine Fourier transforms (power spectra) alongside the corresponding real space reconstructions. The Fourier transform of the original weighted back-projection shows the typical bow-tie shape, reflecting the tomographic missing wedge where projection images from high tilt angles are lacking. The constriction of information to the vertex at the origin implies poor axial resolution in real space, i.e., smeared contrast. The Wiener filter had little effect, but the two iterative deconvolution methods succeeded in “filling in” the missing wedge. Equivalently, the real-space image of the bead is constrained spatially in the axial direction; to the extent that the bead appears as a small Gaussian spot, the 2D Fourier transform of the corresponding plane appears as a broad Gaussian patch. The ER-Decon approach succeeded in restoring spherical symmetry so that all three views appear essentially identical in both real and Fourier space. The dual benefits of deconvolution, restoration of spherical shape and suppression of projected intensity, are seen clearly in an orthoslice view of three perpendicular planes with a gold particle situated at the origin (Fig 2E). Lacking the spurious streaks to guide the eye, the small spherical particle even becomes difficult to discern at the origin. Gold beads are a relatively easy case, however, due to their structural simplicity and strong signal. A more rigorous test of the method would be complex biological structures with realistic low contrast.

We next tested the performance of deconvolution on a cellular tomogram. The bright field recording was used, with contrast inverted so that density appears bright. A central slice through the full reconstructed field of view (∼4×4×0.7 μm) appears in Fig 3A. A pair of mitochondria appear, containing prominent matrix granules of amorphous calcium phosphate (26), as well as a prominent pair of membranous organelles, apparently autolysosomes (34), rough endoplasmic reticulum, and polyribosomes. Microtubules are clearly visible in other planes. We first selected the autolysosomes as an example of low contrast structures. Fig 3B shows side by side views of the reconstruction (low-pass filtered) and the deconvolution as orthoplanes; these are animated in Movie 1 to sweep across the volume. Following deconvolution, internal features of the organelles become clear while distinct structures appear above and below. As a second example we examine the mitochrondria in Fig 3C,D. Here the bright membrane cristae interdigitate among the dense granules. The cristae are clearly resolved in all three axes after deconvolution as seen in Fig 3C. The same features can be found in the filtered reconstruction showing the same slices, Fig 3D, but it would be very difficult to discern them from noise a priori.

**Figure 3.**
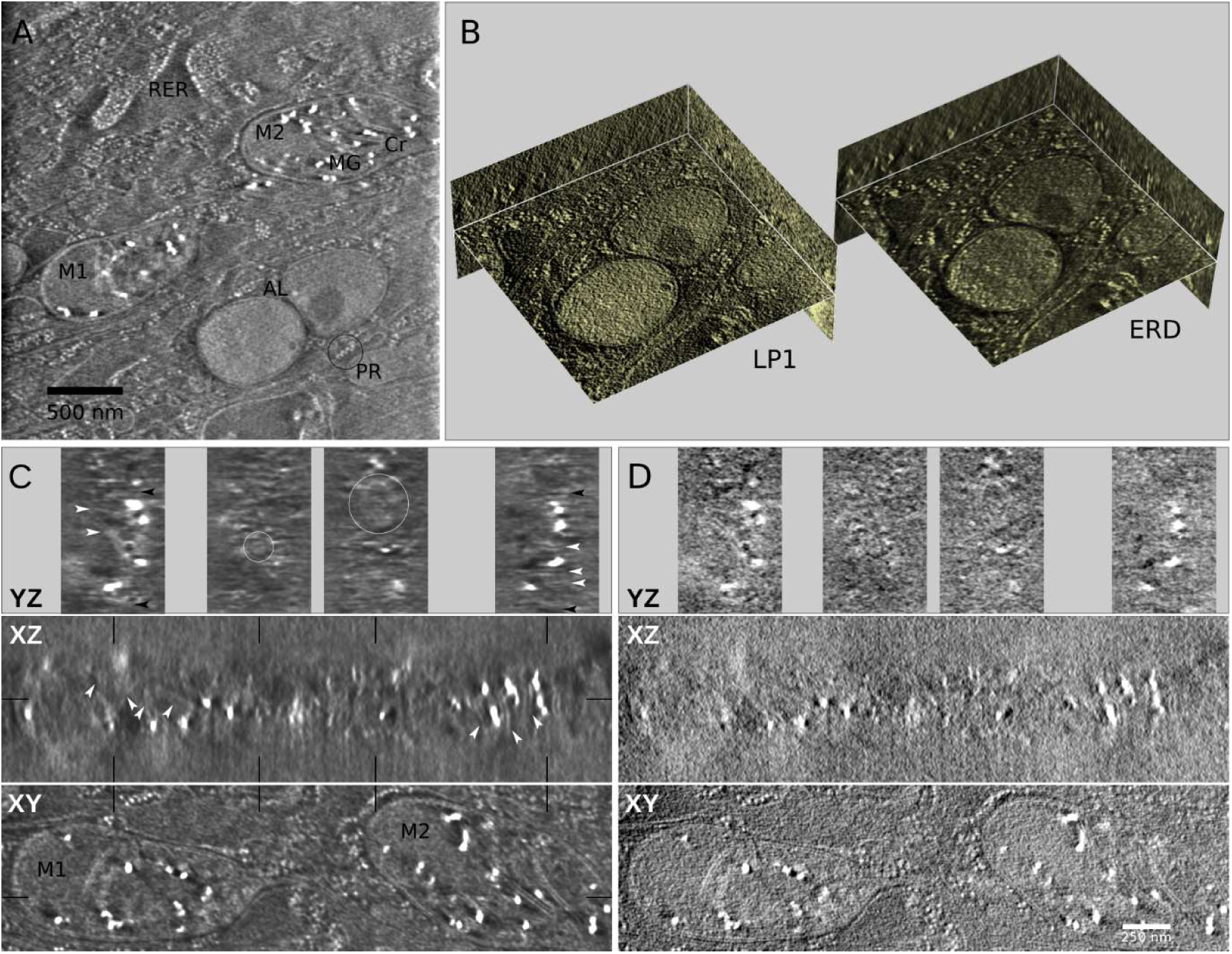
Deconvolution enhances interpretable contrast in whole cell CSTET. A) A single section from the deconvolved volume shows an overview of the cytoplasmic content under investigation. Rough endoplasmic reticulum (RER), polyribosomes (PR), two mitochodria (M1 & M2) with internal cristae (Cr) and calcium phosphate matrix granules (MG), and two putative auto-lysosomal compartments are annotated. B) 3D reconstructions of the AL are shown in orthoplane views after application of a low pass filter (Gaussian, radius = 1 pixel: LP1) and entropy-regularized deconvolution (ERD). The images are opening frames from an animation available in the Supporting Information (Movie 1). C,D) Orthoplane views of the mitochondria with deconvolution and weighted back-projection (filtered as above) respectively. The two panels show identical sections; panel C is annotated. Contrast is enhanced by deconvolution in the XY views, but the most dramatic improvement is seen in XZ and YZ sections. Thin black lines indicate the positions of the sections shown above. Note the cristae (white arrowheads) interdigitated between high contrast matrix granules in M2, the mitochondria boundaries (black arrowheads), the tubular extension emanating from M1 (small white circle), and a cut through the end of M2 where the double membrane can be discerned (large white circle). Similar features are visible in the back-projection but are buried in noise of comparable intensity. Image intensities are scaled linearly based on the volume histogram with a small fraction (0.1 - 0.5%) of voxel values saturated.

Iterative deconvolution typically involves one or more adjustable parameters. In the case of ER Decon II, these are a “non-linearity” in the intensity mapping and a “smoothing” imposed in computing the error function. The latter has a strong qualitative influence, as seen in Fig 4 where its value is scanned across the reasonable range. Within that range it can be tuned to enhance interpretation. Indeed several runs should be performed and compared with each other and with the original dataset.

**Figure 4.**
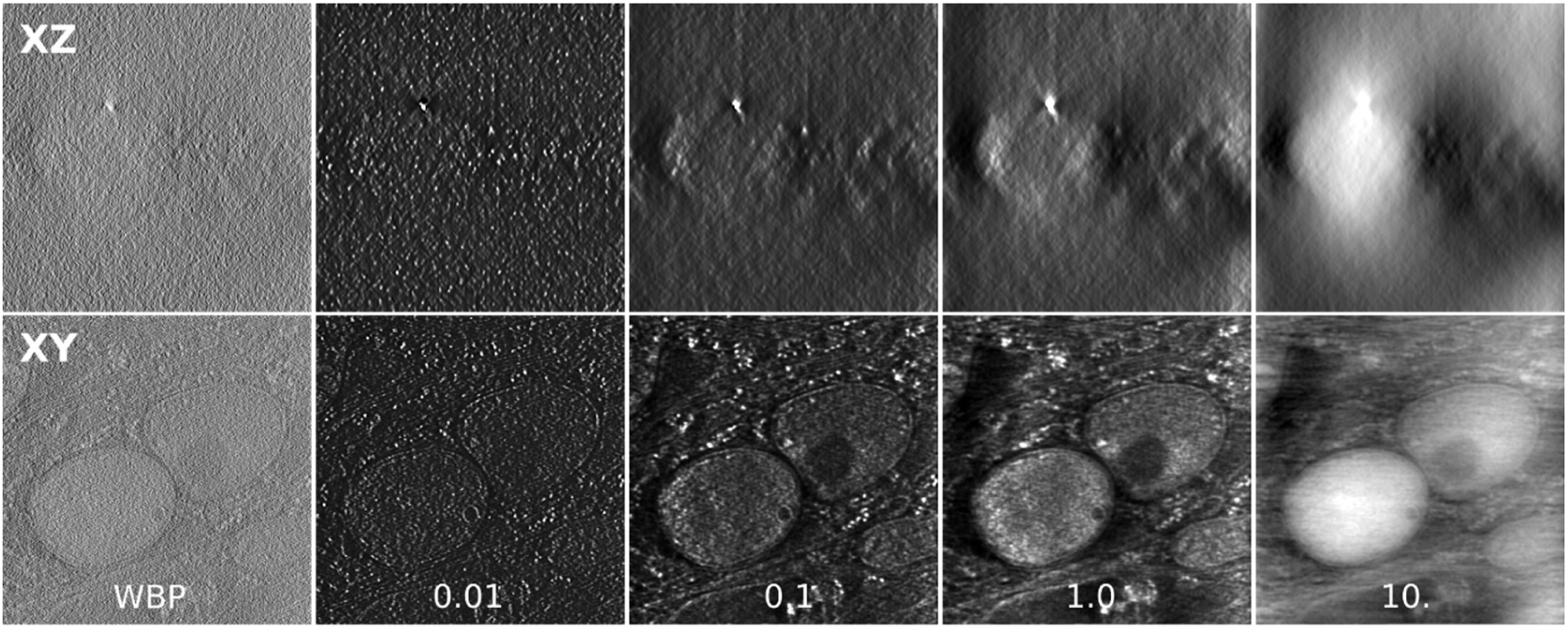
Deconvolution parameters. XY and XZ sections are shown for the weighted back projection (unfiltered) and deconvolution with smoothing values as indicated. See Supporting Fig 3 for location of the chosen sections. Display scaling is linear in all cases with 0.1% of the histogram saturated at high and low values.

Fig 4 shows the effect of the adjustable smoothing parameter on the low contrast data of Fig 3B. As for the gold beads there is a range of suitable smoothing parameters, outside of which obvious artifacts appear. Next we consider mitigation of the missing wedge effect using this realistic dataset (smoothing = 0.1). 2D power spectra (2DPS) were computed slice by slice for the XZ planes and then averaged. The result from the reconstruction (Fig 5A) reveals a set of discrete radial lines perpendicular to each of the projection angles. A similar analysis applied to the deconvolution generates instead a smooth bow-tie shape (Fig 5B). Its limited radius is equivalent to a low-pass filter; the very highest frequencies have been suppressed (comparably to Fig 3). More significantly, the spines partly filled in to produce a continuous distribution. Effectively, it appears that the deconvolution has interpolated across the discrete samplings in Fourier space. Looking carefully, some intensity also appears within the big missing wedge. Possibly the effect of the missing wedge depends on the brightness of the represented features. Intensity histograms appear in Fig 5E. For the original reconstruction, the distribution is Gaussian as expected. After deconvolution, it is strongly skewed to positive values with a long tail to the bright side. Thresholds were then imposed on the dataset and the 2DPS averages recalculated, Fig 5C,D. As seen, the big missing wedge is absent at the highest intensities and appears most prominently in the lowest, where there is anyhow little image detail.

**Figure 5.**
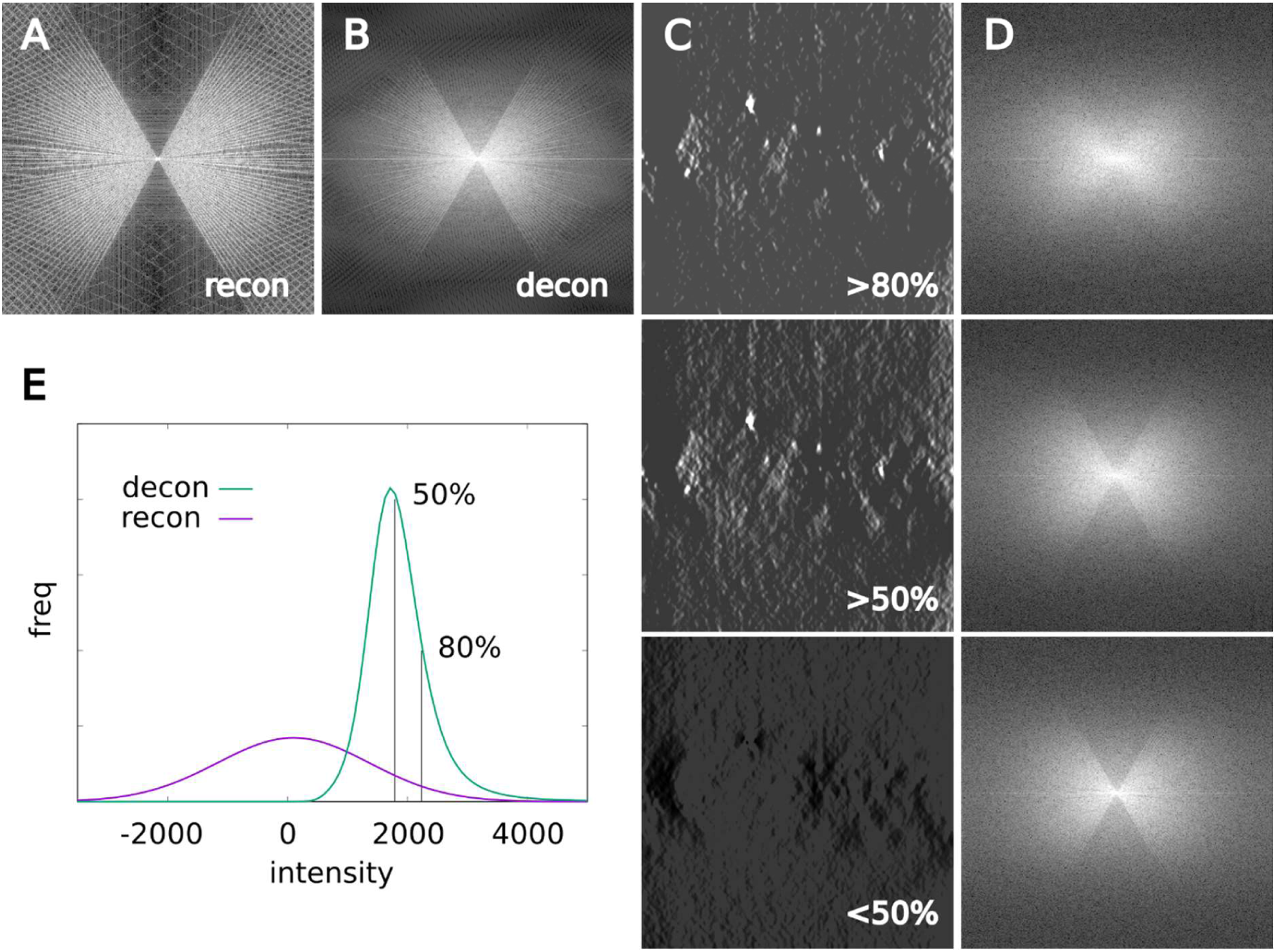
Amelioration of the missing wedge artifact. A,B) Averaged 2D power spectra from XZ planes of the data cubes displayed in Fig 4 for the unfiltered reconstruction and the deconvolution with smoothing = 0.1. C) A central slice of the data thresholded for the lower 50%, upper 50%, and upper 80% of voxel intensities. (The extreme 0.1% were allowed to saturate as in Fig. 4.) D) Averaged 2D power spectra (logarithmic scaling; see Supporting Fig S3 for linear scale) for the corresponding thresholded volumes. E) The volume histogram of the reconstruction reflects the normalization of the raw images prior to back-projection; the histogram of the deconvolution is strongly distorted, with features of interest pushed to the bright tail of the distribution.

As a second example of cellular tomography we chose an area showing the nuclear envelope, with a nuclear bleb, or “polyp”, extending into the cytoplasm as seen in Fig 6. A long, thin mitochondrion containing matrix granules (26) lies adjacent to the outer nuclear envelope at the neck of the extension (Supporting Information Movie 2). Within the nucleus, significant density differences are seen in the chromatin, particularly at the nuclear envelope and inside the polyp. After deconvolution the contrast is greatly enhanced so that these differences become unmistakably clear, yet resolution remains sufficiently high to follow molecular strands in the sparse areas. It is striking that the dense regions take an extended linear form with indication of a twisted structure. Bands can be seen in certain locations along the heavy filaments, whereas the sparsest areas are still filled with thin strands joining bright spots, presumably nucleosomes. The boundaries between dense and sparse chromatin regions are too intricate for a manual segmentation. Instead, we prepared a highly smoothed version of the deconvolution to create a low-resolution map. Application of an isosurface threshold encloses the dense structures, whose diameter is typically 100-200 nm. Interestingly, the dense structures are largely connected in 3D as seen in Fig 6C. This is consistent with the notion of heterochromatin as a macroscopic condensation as opposed to a patchwork of isolated elements (35, 36).

**Figure 6.**
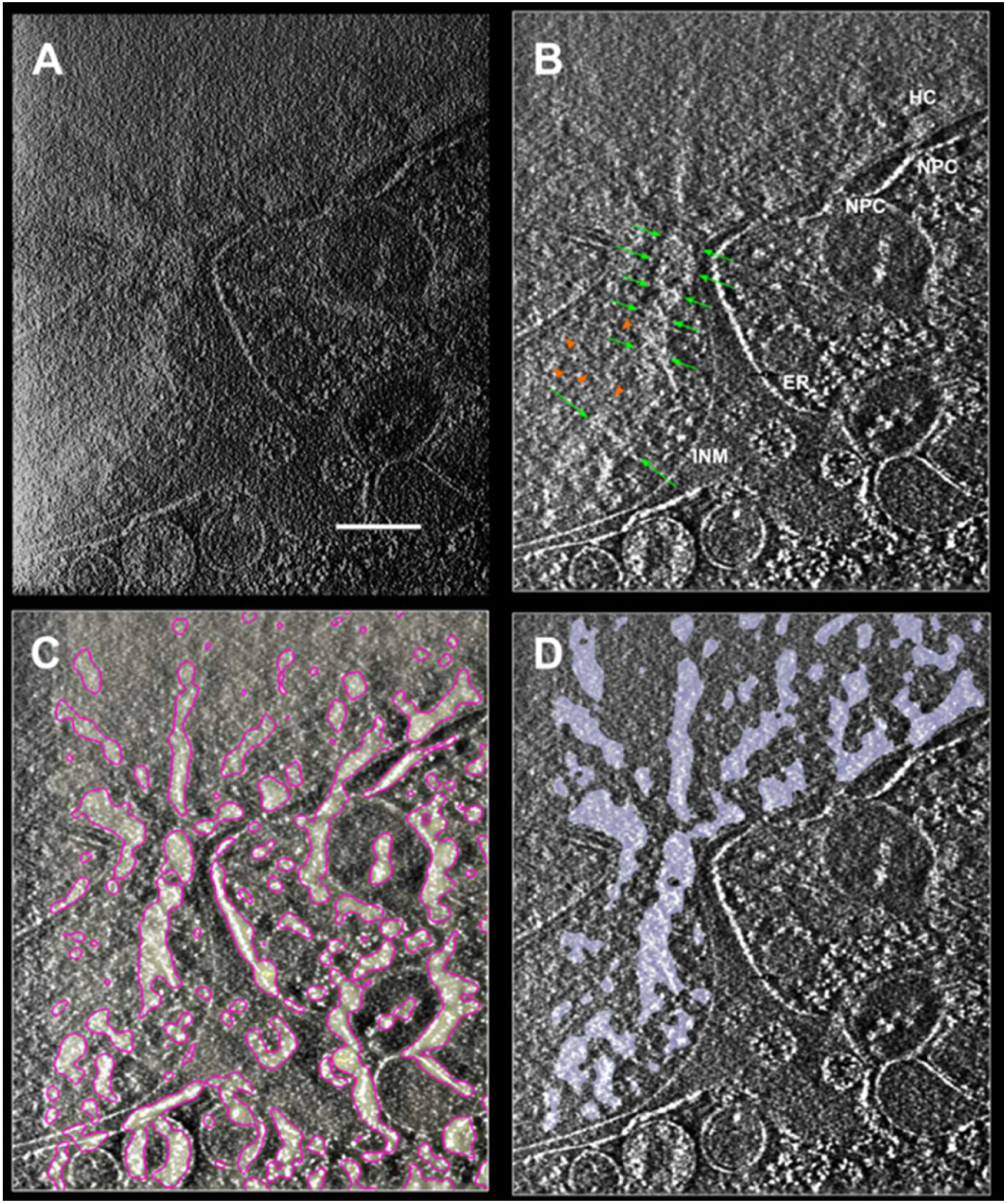
Distinct densities inhomogeneities in interphase chromatin. The nuclear envelope in this fibroblast cell displays a bleb, or “polyp”, into which the chromatin spreads. The outer membrane of the protrusion separates into rough endoplasmic reticulum (ER). The inner nuclear membrane (INM) delimits the chromatin. Nuclear pores (NPC) and perinuclear heterochromatin (HC) are visible near the nuclear envelope at the upper right. The polyp outer membrane makes a junction with a mitochondrion passing underneath the section displayed. See Supporting Information Movie 2 for a scan of the entire volume. A) The raw reconstruction, low pass filtered. B) Deconvolution improves contrast against the background. Within the bounds of the INM, higher density (brighter) regions appear in extended linear structures, both in the perinuclear heterochromatin and interior structures. The dense regions show a twisted shape (green arrows), whereas isolated strands appear in the sparser regions (orange arrowheads). C) A highly smoothed version of the deconvolution follows the denser features. A 3D mask is created by applying an intensity threshold to the smoothed deconvolution, shown here as one section in pink. D) Application of the mask to the data of panel B, limited additionally by the INM, segments the denser chromatin structures (shown in blue) in an unbiased manner. All images present a single section near the center of the reconstructed volume. The pink boundary is displaced by one section for clarity. The total reconstructed thickness near the polyp neck is 1 μm. Scale bar 500 nm.

## Discussion

We have shown that CSTET reconstructions are significantly improved by 3D deconvolution. A previous application used deconvolution to extend the short depth of field in through-focus series of aberration-corrected STEM images (37); this is optically similar to fluorescence imaging with high numerical aperture optics. The present application is to tilt-series tomography, which requires a synthesized 3D PSF as described. Would it be possible to acquire projections from all angles continuously, the back-projection should in principle return the original density. In practice, the set of projections is neither complete nor continuous. Since the resulting artifacts are effectively suppressed by iterative deconvolution, the processing interpolates to some extent across missing data and so can be regarded as a means to reduce exposure of sensitive cryogenic specimens.

Side by side comparison of volumes with and without deconvolution shows that all features present in the deconvolution can be traced to the original reconstruction. The intensity histogram of the deconvolution is strongly distorted, however. Sharp, bright features are promoted to high intensities while the undesirable projection of contrast from other planes is relatively suppressed. This is conceptually similar to the removal of out-of-focus haze by fluorescence deconvolution. The essential insight is that a bona fide bright spot can be distinguished from noise by its 3D structure in the back projection. Deconvolution enhances local intensity to the extent that a match is found with the PSF; this results in suppression of the familiar missing wedge artifacts. Note that post-processing such as deconvolution cannot restore information that was never recorded. It can, however, improve the fidelity of a representation.

A number of different approaches have been taken to address these issues. For example, iterative reconstruction with non-uniform fast Fourier transforms (INFR) can interpolate some information into the missing wedge (38). Filtered iterative reconstruction technique (FIRT) and Iterative Compressed-sensing Optimized Non-uniform fast Fourier transform reconstruction (ICON), impose reasonable assumptions on the smoothness of biological specimens in order to interpolate missing data so as to improve the reconstruction quality (39, 40). Model-Based Iterative Reconstruction (MBIR) incorporates a physical model of the image formation (41). Missing Wedge Restoration based on a Monte Carlo evaluation of randomly introduced noise can extend some information into missing Fourier components based on consistency with the sampled data (42). Finally, inpainting and deep learning approaches begin to emerge for optimization of tomogram reconstruction (43, 44), annotation (45), and particle extraction (46). Deconvolution is a deterministic algorithm that exploits prior knowledge about the 3D representation of a point object. Its utility is well established in fluorescence microscopy, and it could likely be extended or combined with other reconstruction methods so long as the PSF is well defined. The unipolar contrast in STEM, as well as the additive nature of the back-projection, lend themselves particularly well to application of deconvolution in CSTET.

The advantages of CSTET and deconvolution combine to reveal striking structures in chromatin, as seen in Fig 6. We identify the dense features with heterochromatin, e.g., the structures lining the nuclear envelope, and the sparser areas with euchromatin. The distinction between these chromatin forms is ambiguous and depends largely on the assay, e.g., by DNA or nucleosome density, histone modifications, or binding of heterochromatin-associated proteins. There is little evidence to correlate these various descriptions. Strong heterochromatin-euchromatin contrast is of course seen by metal staining in embedded specimens using TEM or FIB-SEM (47). The image contrast depends on differential adsorption of the metal stain, primarily to protein, however, and not on the DNA density directly. A recent breakthrough in osmium staining via singlet oxygen-induced polymerization of diaminobenzidine provides unambiguous labeling of the DNA at the level of nucleosome (48). Still, the use of chemical fixative may influence larger scale structures that lack an internal support. Detection of chromatin density inhomogeneities by cryo-TEM has been challenging due mainly to constraints of phase contrast, including specimen thickness and the overall flat image texture. (The latter is a general property of defocus phase contrast, wherein the contrast transfer function goes to zero at low spatial frequencies. Phase plate technology restores contrast at low spatial frequency, like STEM, but the specimen thickness is still limited by loss of phase coherence.) STEM imaging relaxes these constraints, so that highly complex features are easily segmented on the basis of density at low resolution. Extended chromatin structures with diameter on the order of 100 nm and an overall twisted appearance suggest a mesoscale organization within the chromatin territories of the interphase nucleus. Thus cryo-STEM tomography in combination with deconvolution provides important new opportunities for 3D imaging of fully hydrated, intact cells.

## Acknowledgements

The authors are grateful to Muthuvel Arigovindan, Shahar Seifer, Charles Kervrann, David Agard, and David DeRosier for enlightening discussions. This work was supported in part by a grant from the Israel Science Foundation. ME is incumbent of the Sam and Ayala Zacks Professorial Chair and head of the Irving and Cherna Moskowitz Center for Nano and Bio-Nano Imaging. The lab has benefited from the historical generosity of the Harold Perlman family.

PRIISM software and ER-Decon II are available for academic use upon request from John Sedat or David Agard, UCSF.

