## Supplementary Information for "3D Deconvolution Processing for STEM Cryo-Tomography"

**Supporting information**

Includes: detailed protocol for deconvolution

Supporting figures S1-S3

Movies 1 & 2, related to figures 3 and 6.

**Protocol for deconvolution**

Following is a recommended protocol for 3D deconvolution of tomographic datasets. The procedure consists of 3 basic steps: construction of a point spread function (PSF), conversion of the PSF to an optical transfer function (OTF), and implementation of the deconvolution.

As in fluorescence microscopy, the PSF is ideally the shape of the 3D image produced by a point source. It can be approximated by a “theoretical” PSF, which for fluorescence microscopy is the focused cone produced by the objective, taking into account the emission wavelength. Since STEM contrast is based on scattering, and on assumption of a linear relation between the integrated scattering cross-section and scattering away from the bright field detector, we can approximate the PSF by the illumination profile. While electron optics suffer inherently from spherical aberration, tomography data were collected here with a very small semi-convergence angle (I.e., numerical aperture), primarily in order to extend the depth of focus. As a result, we can approximate the illumination profile as a diffraction-limited wave, again using tools for optical imaging. Each image in the tilt series is recorded with the same PSF, but inclined to a different angle. In the back-projection their contributions are summed, so as a 3D PSF we use the sum of the diffraction-limited wave profiles, tilted to the same angles used in back-projection.

**PSF:** Ideally the PSF is created at the same size as the volume to be deconvolved. A single beam is first prepared using the Diffraction_PSF_3D plugin in ImageJ, and then tilted and summed using a custom ImageJ macro as shown. Ensure first that the PSF window is renamed to “PSF”.

```
selectWindow("PSF");
w=getWidth();
h=getHeight();
z=nSlices;
run("Reslice [/]...", "output=1.000 start=Top avoid");

selectWindow("PSF");
close();
newImage("Untitled", "32-bit black", w,h,z);
run("Reslice [/]...", "output=1.000 start=Top avoid");

pathfile=File.openDialog("Choose the .tlt file:");
filestring=File.openAsString(pathfile);
lines=split(filestring,"\n");
n_lines=lengthOf(lines);
Ar=newArray(n_lines);

for(n=0; n<n_lines;n++)
  {
Ar[n]= lines[n];
  }

for(j=0;j<Ar.length;j++)
 {
theta_i=Ar[j];
selectWindow("Reslice of PSF");
run("Duplicate...", "duplicate");
run("Rotate... ", "angle=theta_i grid=1 interpolation=Bilinear stack");
imageCalculator("Add 32-bit stack", "Reslice of Untitled","Reslice of PSF-1");
print("rotated by", theta_i);
selectWindow("Reslice of PSF-1");
close();
  }
```

**OTF:** The STEM PSF is converted to OTF by a 3D Fourier transform (power spectrum). This is done conveniently in Priism software with the 3DFT tool. First, the orientation should be flipped so that the Z axis corresponds to the beam propagation, with the tilt axis vertical to match the orientation of IMOD aligned series. Using the Flip tool (DataViews > Rotation > Flip), X-Z Sections:

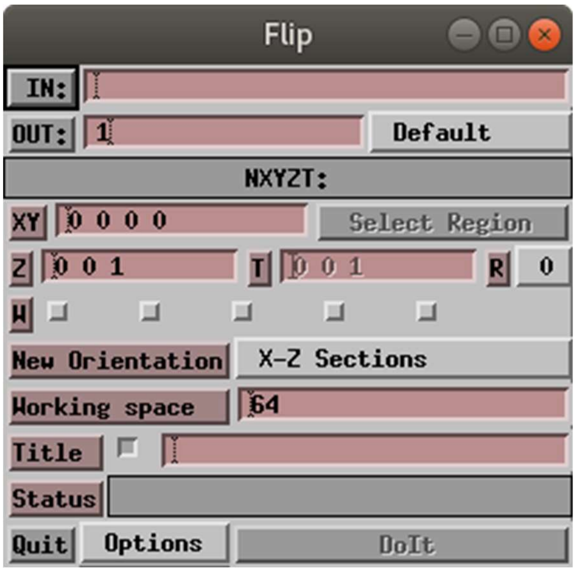


At the end, Do It.

The pixel sampling should be chosen to match that of the volume data. In Priism, EditHeader (Dataviews > DataInfo > EditHeader), Pixel spacing:

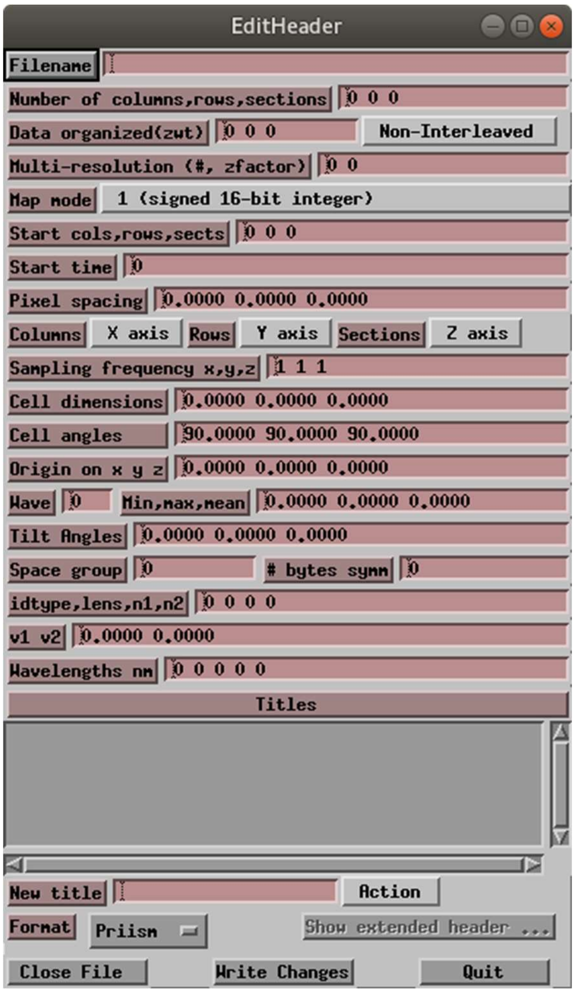


Take care to match the header info of the reconstructed volume, and not to mix units (e.g., Angstroms with nm). At the end, Write Changes.

Finally, create the OTF (Processing > Fourier Tools > 3D Transform).

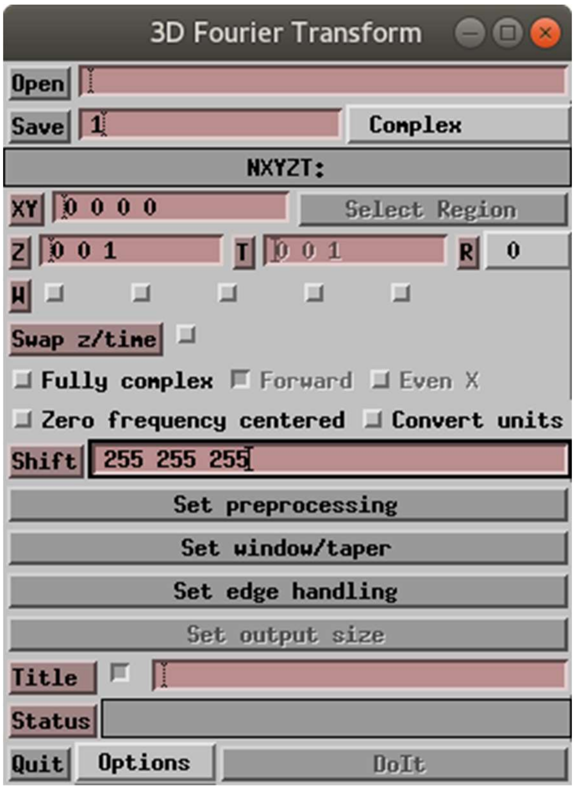


Be sure to uncheck Convert units (as displayed) and to edit the Shift entry to match the center of the PSF volume (as displayed for a volume 512^3).

At the end, Do It.

**Decon:** Deconvolution is performed with the core2_decon program, which can be accessed by GUI from the Priism main menu (Processing > OM Processing > ER-Decon II).

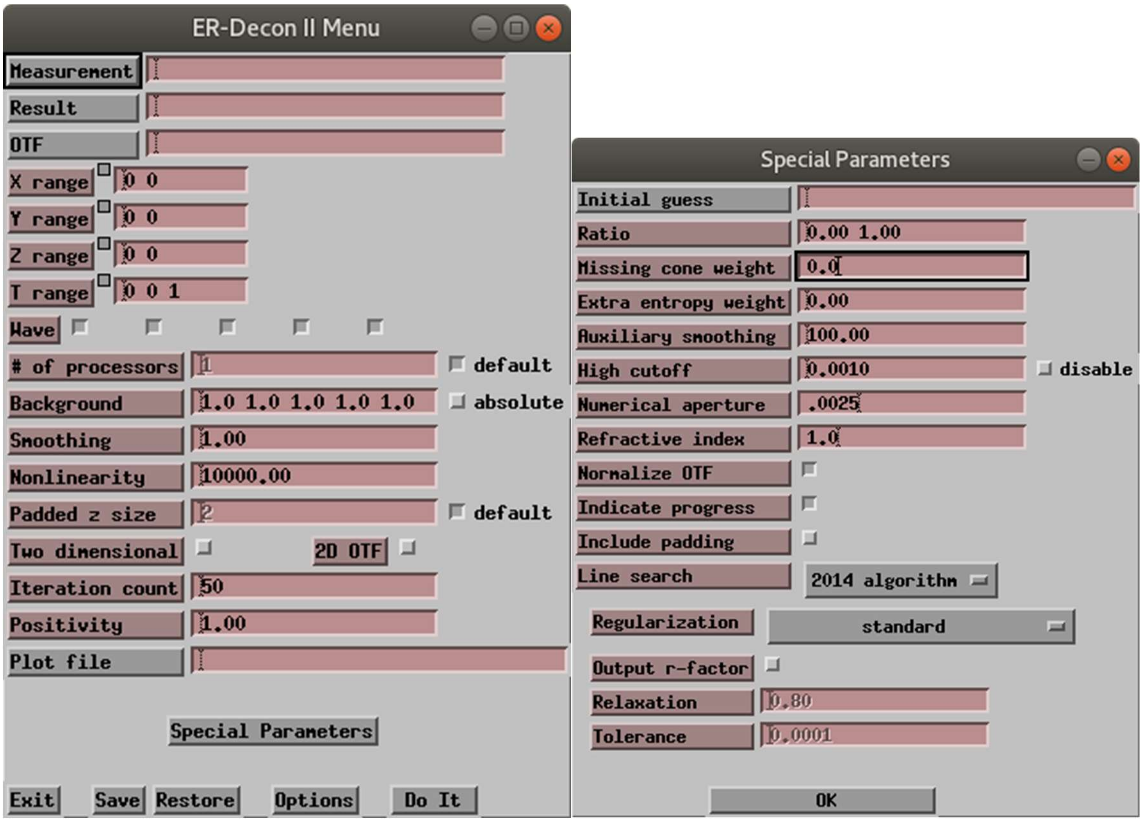


Measurement specifies the reconstructed volume, Result the output, and OTF file. Essential parameters are Smoothing, Nonlinearity, and Iteration count. In Special Parameters, the Missing cone weight must be set to 0. Numerical aperture and Refractive index should be ignored, but are set to match the PSF generation. Indicate progress is useful in the log file.

At the end, Do It.

In addition to the GUI, Priism functions can be executed from the command line. This is especially useful for core2_decon, which can take easily several hours and is conveniently run by batch submission on a strong server. Such command files can be generated from the GUI using the Options button, filling in filenames for com and log, and selecting “Skip execution”.

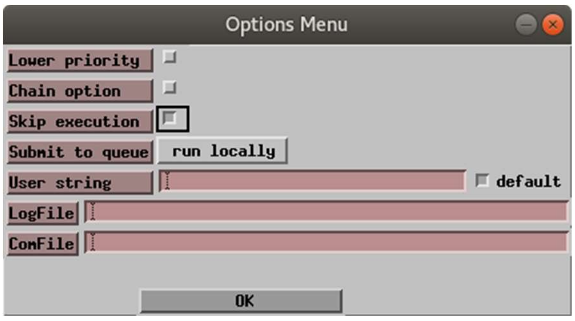


An example for the 3D Fourier transform is as follows, where parameters in brackets < > should be filled in appropriately by the GUI, or else manually to match the data:

```
#!/bin/sh
echo Running on `hostname` >'3DFT.log'
#Setting run time environment...
{ test -r '<path to Priism>/Priism_setup.sh' && . '<path to Priism>/Priism_setup.sh' ; } || exit 1
(time FTransform3D \
 '<PSF filename>.mrc' \
 '<OTF filename>.otf' -x1=<0:511> -y1=<0:511> -z1=<0:511:1> -w1=0 -t1=0:0:1 \
 -res1=0 -mode1=complex -title='' -real_complex -same_units \
 -shift=<255:255:255> -xpad=default -xpad_value=0:0 -ypad=default \
 -ypad_value=0:0 -zpad=default -zpad_value=0:0 -3d=z ) >> '3DFT.log' 2>&1
```

For core2_decon it is useful to have parameters appear in the filename. This can be done by editing the GUI-generated script, for example:

```
#!/bin/sh
## usage: ./core2-run <input filename>.rec <smoothing> <non-linearity> <OTF filename> <number of cycles>
## note: The input file is specified without the .rec or .mrc extension. The output decon file will have the standard .mrc extension
## z padding is hard wired here. Normally the PSF should be built at the same size as the processed volume.
echo Running on `hostname` > `pwd`/core2_decon-$1_s$2_nl$3_nc$5.log
#Setting run time environment...
{ test -r '<path to Priism>/Priism_setup.sh' && . '<path to Priism>/Priism_setup.sh' ; } || exit 1
#command file for core2_decon
( time core2_decon $1.rec decon-s$2_nl$3_nc$5-$1.mrc $4 \
-alpha=$3 -lamratio=0:1 -lamf=$2 -lampc=0 -lampos=1 -lamsmooth=100 \
-laml2=0 -cuth=0.001 -ncycl=$5 -nzpad=512\
-omega=0.8 -sub=1:1:1:1:1 -tol=0.0001 -np=4 -oplotfile="plotfile-$1_s$2_nl$3_nc$5" \
-linesearch="2014" -regtype="ma" -logging=progress ) \
>>core2_decon-$1_s$2_nl$3_nc$5.log 2>&1
```

Choose other parameters at will, for example padding in z (-nzpad) or number of processors used for execution (-np), according to the Priism help pages.

**Evaluation of the deconvolution performance**

We find several checks to be useful in evaluating the quality or reliability of a deconvolution.

The most obvious criterion for any image processing procedure is that it should not invent features that were not present in the original data. The best test is side by side visual comparison, in combination with a simpler noise-reduction algorithm such as low-pass filtering as shown in Fig 3B&C. If successful, the deconvolution provides an interpretable contrast with less severe loss of resolution as shown. We have never observed features invented de novo by the deconvolution. As demonstrated in Fig 4, inappropriate parameter selection produces contrast that is not more readily interpretable than the original.

The major pitfall to successful application is a poor match of the theoretical PSF to the reconstruction. The PSF is generated by a pure rotation around a fixed axis. Tomographic reconstruction requires precise alignment of the tilted projection images. The ideal PSF will not match a reconstruction generated with an imperfect alignment. The best test for alignment is a close inspection of the spines radiating from sharp points such as fiducial markers in the XZ plane; these should meet in a point as shown in Fig 2 and S1A. This is in any case a standard quality control in tomography. In a successful deconvolution the fan shape should be reduced to a point, the diameter of which is influenced by the original electron probe diameter and pixel sampling in addition to parameters of the processing.

It is also crucial that the PSF should be constructed from the set of angles actually used in the back-projection, rather than the nominal tilt angles applied at the microscope. (In the IMOD workflow this corresponds to the .tlt file, rather than the .rawtlt.) Angles corresponding to any tilt images left out from the reconstruction should be removed when constructing the PSF as well. On the other hand, the detailed shape of the diffraction-limited illumination profile had a surprisingly weak influence. This is in contrast to optical microscopy and can be understood in terms of the very small convergence angles involved (i.e., low numerical aperture) in collection of each individual tilt image. Even a mismatch in the theoretical probe diameter or pixel spacing has an effect that is noticeable only at the highest spatial frequencies, similar to a low-pass filter or altered smoothing parameter. Care should be taken in interpreting widths of lines or diameters of point objects when these are close to the width of the diffraction limited probe, but otherwise the PSF construction is quite forgiving provided that the projection angles are correct.

Another possibility for mismatch comes into play with very thick specimens, where the transmitted beam is broadened by multiple scattering. Beam broadening is recognized as a “top-bottom” effect in STEM, with resolution deteriorating with depth in the specimen. This problem is more severe with plastic embedded sections than in CSTET. Still, we expect and observe that deconvolution loses effectiveness for samples much thicker than two elastic mean free paths, i.e., roughly 1.2 microns.

Finally, it is important to keep in mind that the flat contrast of the original STEM reconstruction is on the whole a realistic representation of material contrast based on scattering cross-sections. Its corruption by the structural noise sources inherent in practical tilt tomography prevent a stretch in contrast to assist in interpretation, particularly in the axial direction. Care should be taken in interpreting intensity values quantitatively after non-linear deconvolution, or indeed any iterative non-linear processing, because of the distortion of the intensity histogram. Useful as it is for visualization, this distortion would need to be calibrated for meaningful quantitative interpretation.

Supporting Figures

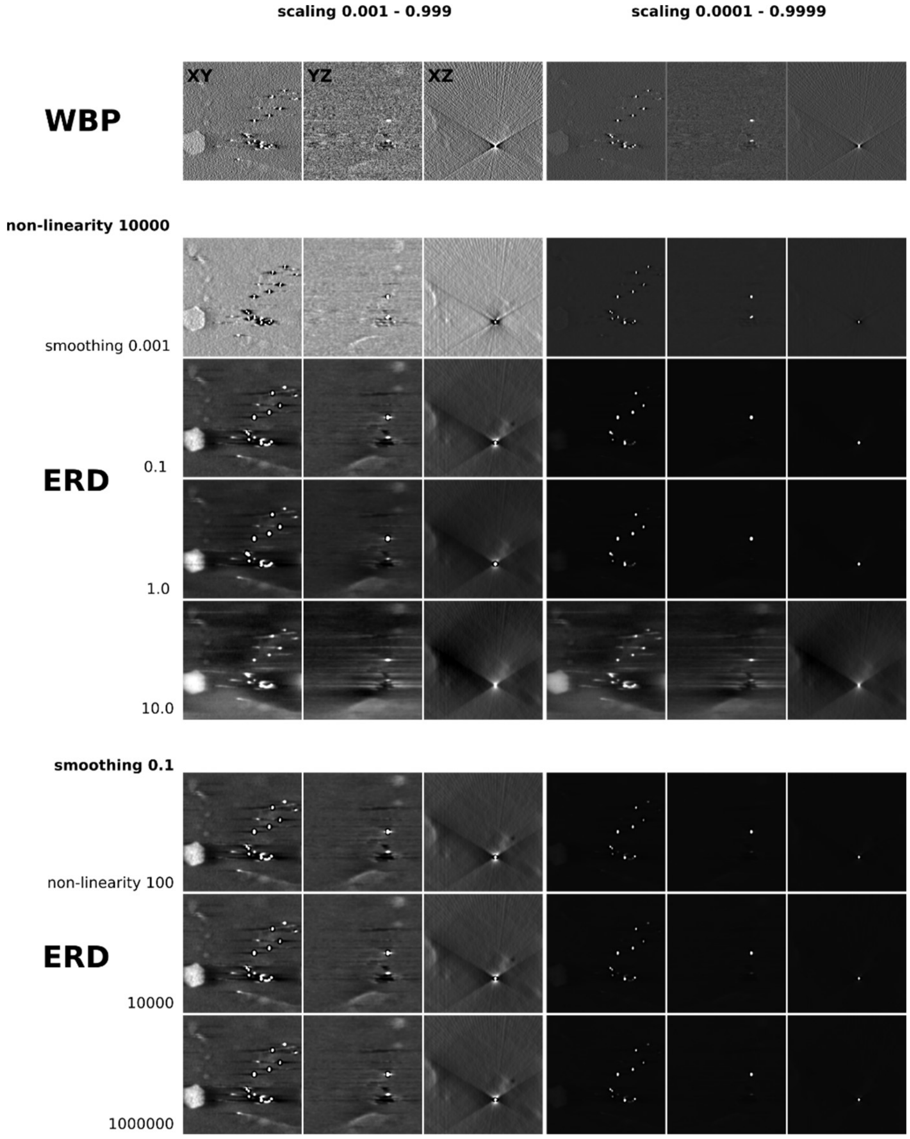


Figure S1. A scan of deconvolution parameters using colloidal gold particles. XY, YZ, and XZ views of the original weighted back-projection (WBP) data appear at the top. Displayed intensity is set to encompass the entire volume histogram except for the extreme 0.1% (left column) or 0.01% (right column) of values. Entropy-regularized deconvolution uses two adjustable parameters: a smoothing and a non-linearity. For fixed non-linearity (ERD, central rows) the smoothing can create strong artifacts when set above or below the suitable range. Gold beads are an easy case in the sense that an intensity threshold can be chosen to hide the faults; this is of course not possible for realistic data with lower contrast. For a fixed and suitable smoothing parameter (ERD, lower rows) the non-linearity has very little influence when the contrast is scaled using the histogram, as seen.

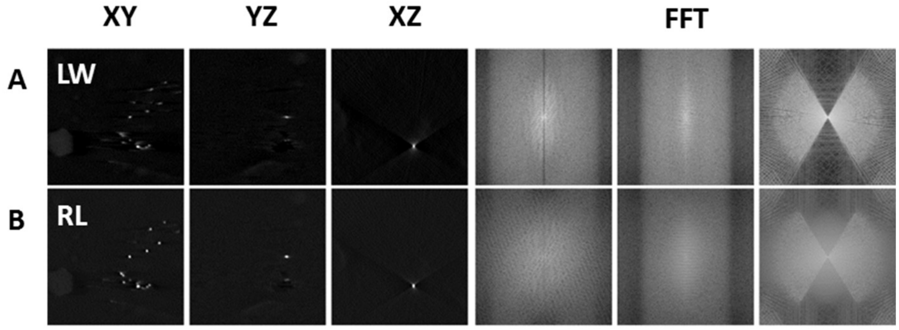


Figure S2. Additional implementations of iterative deconvolution. A) Landweber (LW) and B) Richardson-Lucy (RL) algorithms available in DeconvolutionLab2 . XY, YZ, and XZ slices appear on the left, Fast Fourier Transforms of the same on the right.

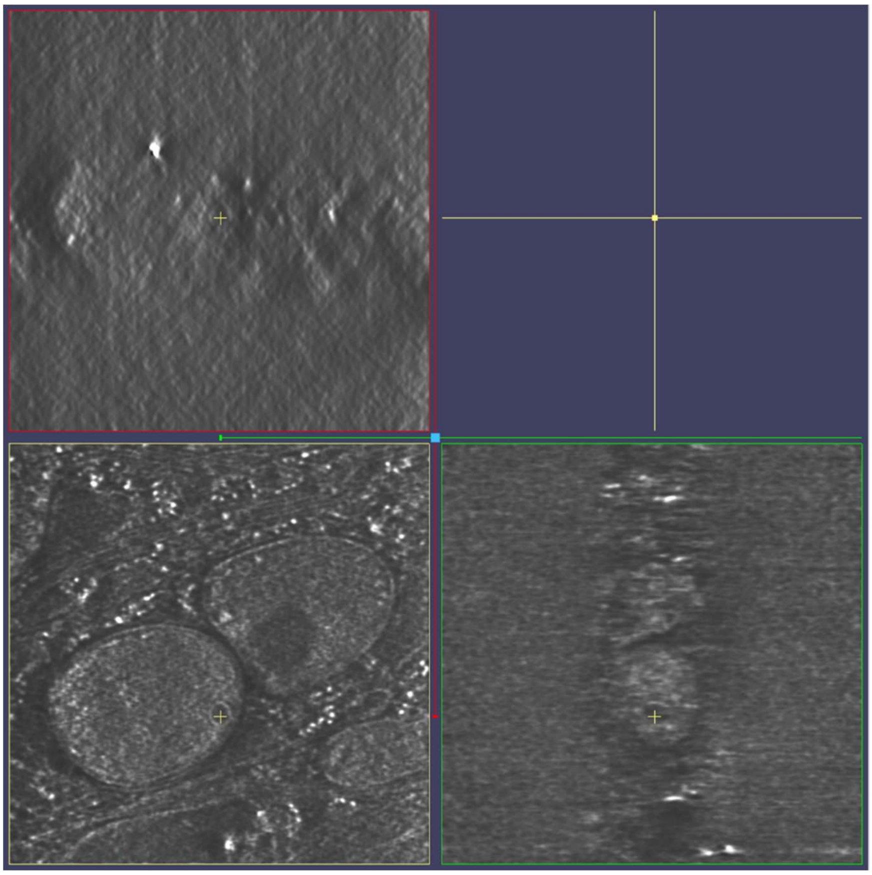

Figure S3. Location of the sections shown in Fig 4 is indicated by the yellow crosshair.

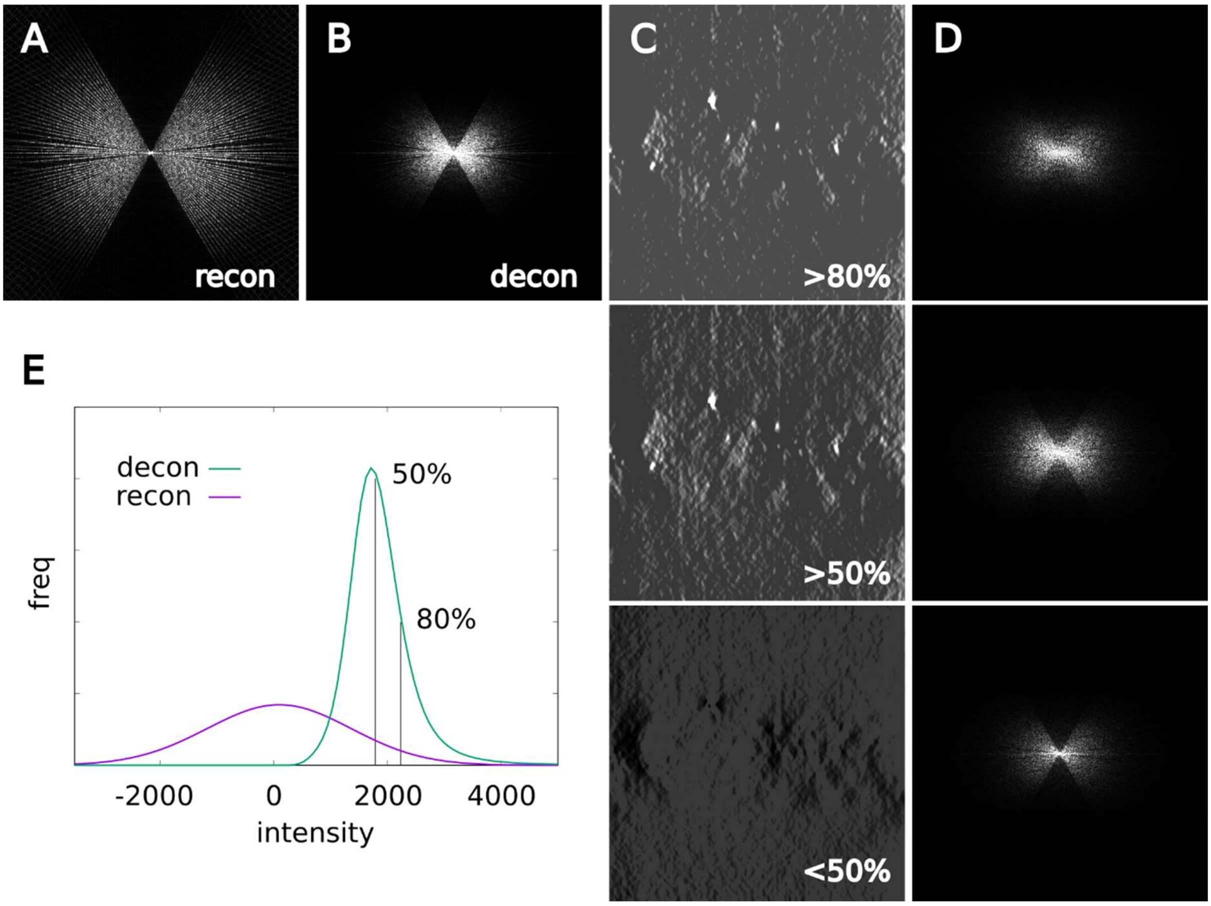


Figure S4. Same as Fig 5 with power spectra shown in logarithmic rather than linear scaling.

Movie 1. Animation of the side by side orthoplane representation of the reconstruction on the left, (Gaussian) low-pass filtered to improve visibility, and the deconvolution on the right as presented in Figure 3.

Movie 2. Segmentation of chromatin to distinct dense and sparse regions. An over-smoothed deconvolution is used as a mask to follow the intricate boundary.
